# Dietary and pharmacological induction of serine synthesis genes

**DOI:** 10.1101/2020.06.15.151860

**Authors:** Alexei Vazquez

**Affiliations:** Cancer Research UK Beatson Institute, Glasgow, UK; Institute of Cancer Sciences, University of Glasgow, Glasgow, UK

## Abstract

There is an increasing interest in the pathway of L-serine synthesis and its. Although L-serine and downstream products can be obtained from the diet, serine deficiency has been documented in neurological disorders, macular degeneration and aging. This evidence calls for strategies to induce serine synthesis. Here I address this problem taking advantage of the wealth of data deposited in the gene expression omnibus database. I uncover that low protein and ketogenic diets increase the expression of serine synthesis genes in the liver and the brain relative to control diets. I discover oestrogen medications, the antifolate methotrexate and serine synthesis inhibitors as classes of compounds inducing the expression of serine synthesis genes in the liver. Future work is required to investigate the use of these interventions for the management of serine deficiency disorders.

## Introduction

L-Serine is a building block of proteins and phospholipids (Fig. 1). L-serine is converted to D-serine in the brain by serine racemase. Serine is also broken down to glycine and formate, providing additional building blocks for the synthesis of purines and glutathione. D-serine and glycine are co-agonists of the N-methyl-D-aspartate (NMDA) receptor together with glutamate, linking serine metabolism to neurological signalling. L-Serine is a nonessential amino acid and mammalians organisms satisfy their serine demand from dietary protein, the serine synthesis from glycine and formate and the serine synthesis from the glycolytic intermediate 3-phosphoglycerate (Fig. 1). Deficiencies in these supply routes may compromise the physiological functions dependent on serine.

**Figure 1.**
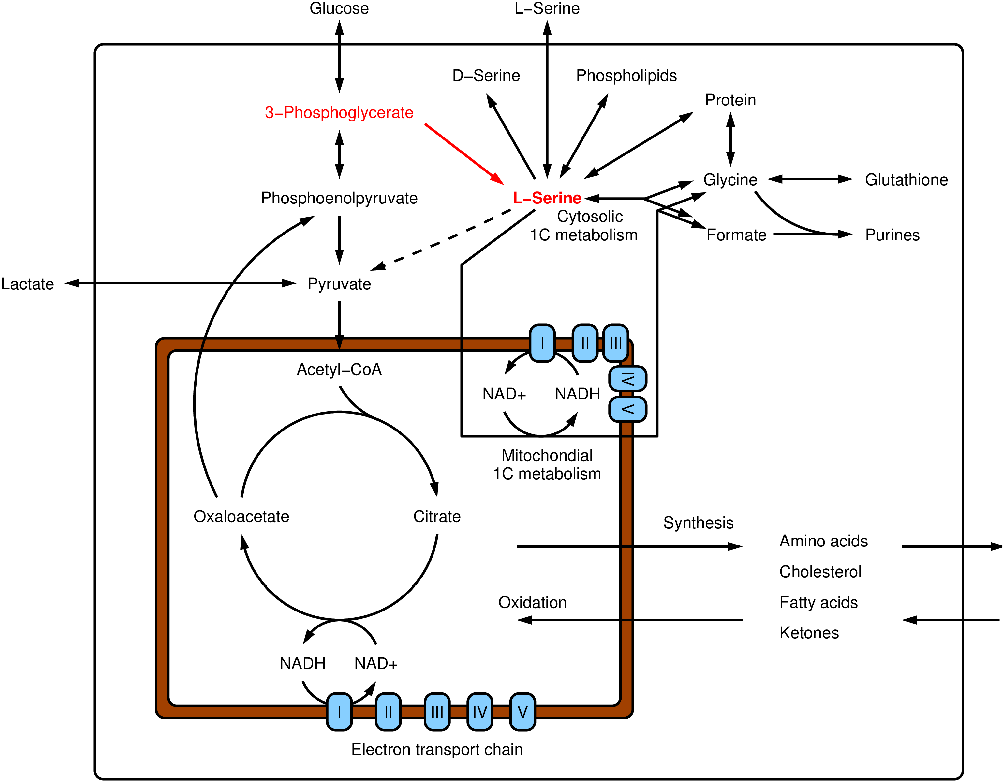
Serine metabolism. Schematic representation of serine metabolism and its interaction with central metabolism. 1C stands for one-carbon.

Serine deficiency has been implicated in physiological disorders in humans and animal models. Inborn errors of serine biosynthesis cause neurological disorders ^1^. Low blood serine is a risk factor for macular telangiectasia type 2 and peripheral neuropathy ^2^. Defective metabolism of serine to glycine and formate impairs the activation of naïve T cells in aging mice ^3^. Homozygous deletion the gene coding for 3-phosphoglycerate dehydrogenase (*Phgdh*), the first enzyme of serine synthesis, causes neurodevelopmental abnormalities and it is embryonic lethal ^4^ in mice.

The manifestation of serine deficiency in the human population call for strategies to increase the availability of serine. Dietary serine supplementation seems the obvious strategy ^5^. However, for a number of reasons, serine supplementation is not an effective approach in general ^5^. Dietary serine is rapidly incorporated into proteins or broken down to glycine and formate ^6^. Mammalian organisms do not have a store for amino acids other than protein ^7^, but proteins are made to perform specific functions and they are degraded as a last resort.

The induction of serine synthesis from the glycolytic intermediate 3-phosphoglycerate is potentially a longer lasting intervention. Here I investigate different strategies to induce the expression of genes encoding for the serine synthesis enzymes.

## Materials and Methods

I have previously introduced a gene expression analysis to identify candidate compounds inducing a specified gene signature in a specified tissue ^8^. For sake of self-consistency, I transcribe sections of that methodology here.

### Gene signatures

The gene signature for serine synthesis contains the genes *Phgdh*, *Psat1* and *Psph*, encoding for the enzymes 3-phosphoglycerate dehydrogenase, phosphoserine transaminase and phosphoserine phosphatase, respectively. The remaining gene signatures were obtained from public repositories or literature reports and they are listed in the Supplementary Table 1.

### Gene expression profiles

The gene expression profiles were downloaded from Gene Expression Omnibus. The accession numbers are GSE57800 (organism: Sprague Dawley rat, gender: male, tissue: heart, control: water vehicle, time: 3 days), GSE57805 (organism: primary Sprague Dawley rat hepatocytes, control: DMSO vehicle, time: 1 day), GSE57811 (organism: Sprague Dawley rat, gender: male, tissue: kidney, control: water vehicle, time: 3 days), GSE57815 (organism: Sprague Dawley rat, gender: male, tissue: liver, control: water vehicle, time: 0.25 days) and GSE57816 (organism: Sprague Dawley rat, gender: male, tissue: thigh muscle, control: water vehicle, time: 3 days) for the pharmacological interventions; GSE144207 (organism: brown rat, gender: male, tissue: liver, control: control diet, time: 4 weeks) for the low protein diet, GSE7699 (organism: C57BL/6 mice, gender: male, tissue: liver, control: chow diet, time: 6 weeks) and GSE115342 (organism: B6.Cg-Lepob/J mice, gender: female, tissue: cerebral cortex, control: regular chow, time: 7 weeks) for the ketogenic diet; GSE51885 (organism: C57BL/6 mice, gender: male, tissue: liver, control: current LAR diet, time: 3 weeks) for the mixed diets and GSE120347 (organism: human neuroblastoma cell line BE(2)-C, control: DMSO, time: 2 days) for the PHGDH inhibitor. In all cases gene expression was quantified using microarrays and the RMA signal (in log_2_ scale) was used as the gene expression readout. For each dataset, the average of the log_2_ expression across controls was subtracted from all samples.

### Gene set enrichment analysis

The significant induction or repression of a given gene signature on a given sample was quantified using Gene Set Enrichment Analysis (GSEA) ^9^ as previously described ^10^. GSEA results in a positive and a negative score quantifying the induction or repression of the gene signature, together with their associated statistical significance (here 100,000 permutations of the genes assignments to probes). A sample was defined positive (red) for a signature whenever it manifested a significant positive score (statistical significance ≤0.05), negative (blue) whenever it manifested a significant negative score (statistical significance ≤0.05) and no significant change (black) otherwise.

### Positive hits

In the input dataset there are multiple compounds tested under multiple conditions (dose and duration of treatment), and each compound/condition was tested in triplicates. A compound tested at a specific condition was deemed positive if it significantly induced the serine synthesis gene signature in the 3 replicates tested. A compound was deemed a hit if it has a significant enrichment of positives across the conditions where it was tested, given the total number of conditions tested and the total number of conditions scoring positive in the whole dataset, using a hypergeometric distribution (Supplementary Table 2).

## Results

### Dietary interventions

The synthesis of serine from 3-phosphoglycerate proceeds through three enzymatic steps catalysed by 3-phosphoglycerate dehydrogenase (PHGDH), phosphoserine transaminase (PSAT1) and phosphoserine phosphatase (PSPH). The genes encoding for these enzymes are under the transcriptional regulation of ATF4, the master regulator of the amino acid stress response ^11^. Experiments with cell cultures demonstrate the induction of the ATF4 transcriptional activity upon deprivation of any of the 21 proteinogenic amino acids ^12^. It is worth asking if the same is true in the context of whole-body metabolism. To address this question, I took advantage of a study reporting liver gene expression profiles of mice subject to a low protein diet (GSE144207, unpublished). As a readout I will use gene signatures for the annotated targets of ATF4, the genes encoding for the serine synthesis enzymes, and other gene signatures relevant to this study.

The low protein diet induces the gene signature for ATF4 targets and serine synthesis (Fig. 2A). There is also a distinctive downregulation of a gene signature for cholesterol synthesis (Fig. 2A), although the exact mechanism behind this observation is unclear. The low protein diet induces the gene signatures of mitochondrial serine one-carbon metabolism (Fig. 2A, 1C mitochondria). The genes in this pathway are also ATF4 targets. This data indicate that a low protein diet activates the ATF4 transcriptional activity in the liver and increases the expression of serine synthesis genes relative to a control diet.

**Figure 2.**
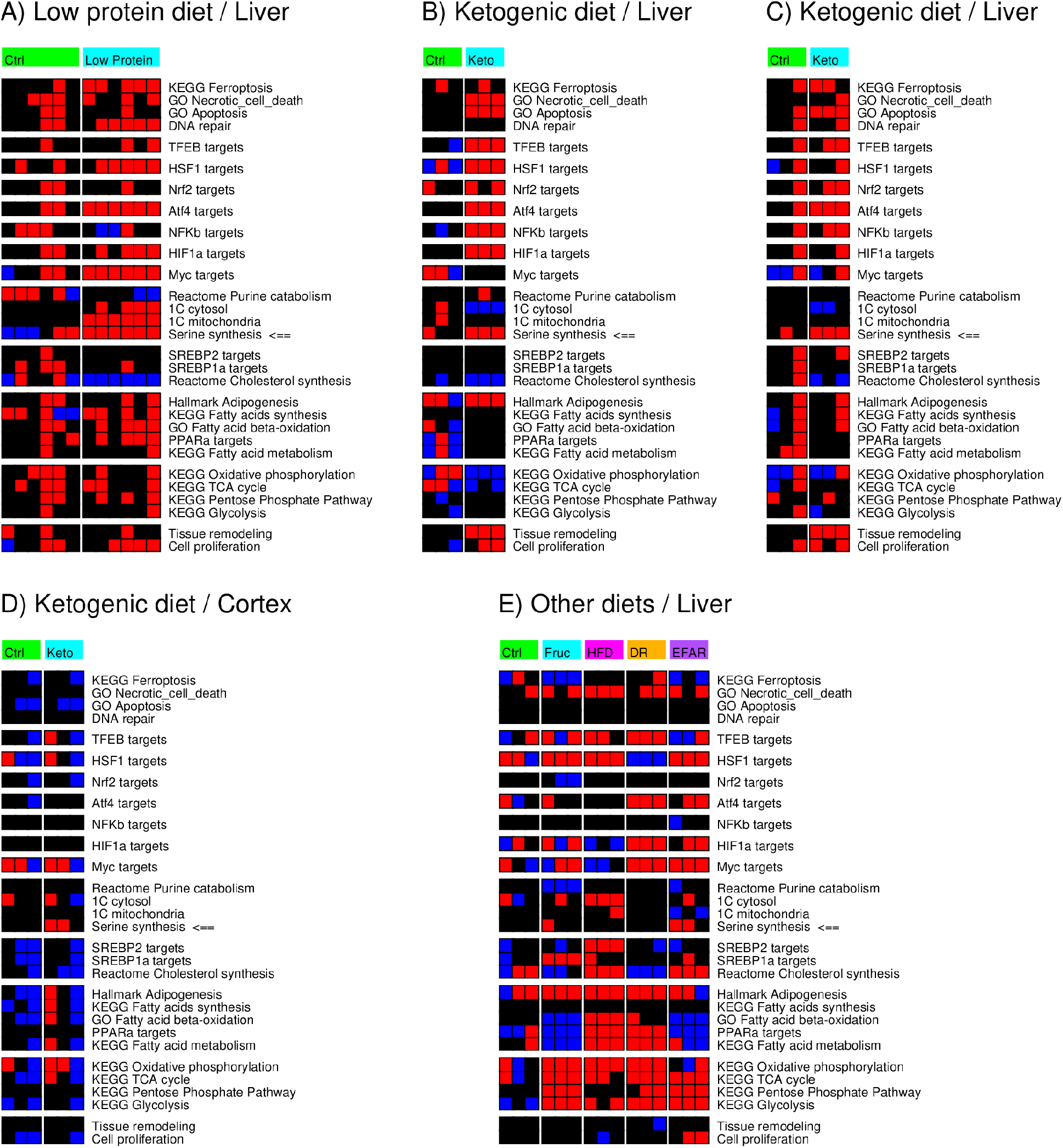
Impact of dietary interventions on the expression of serine synthesis genes. A) Liver gene signatures of mice subject to a low protein diet. B) Liver gene signatures in rats subject to a ketogenic diet. C) Liver and D) cerebral cortex gene signatures in another cohort of mice subject to a ketogenic diet. E) Liver gene signatures in mice subject to different diets. Columns represent samples clustered by intervention and rows represent gene signatures. <⩵ Points to the serine synthesis gene signature. Red indicates a significant gene signature induction relative to the average in controls, blue a significant repression and black no significant change. Keto: ketogenic diet, Fruc: high fructose diet, HFD: high fat diet, DR: dietary restriction, EFAR: essential fatty acids restriction.

As a second dietary intervention I consider a report of liver gene expression profiles of mice subject to a ketogenic diet (GSE7699 ^13^). A ketogenic diet shifts the calorie intake to dietary fat and away from carbohydrates, while maintaining the same protein intake. At first one may think that a ketogenic diet should have no significant alteration on amino acids metabolism. However, we should bear in mind that nonessential amino acids can be made from glucose and a reduction in the availability of glucose may impinge on the availability of non-essential amino acids. This hypothesis is proven to be true. The ketogenic diet induces both the gene signatures for ATF4 targets and serine synthesis (Fig. 2B), effectively acting as the low protein diet. This observation by itself challenges the common believe that sugars are empty calories. The induction of the amino acid stress response by a diet with a low sugar intake is the empirical evidence that sugars contribute to amino acid metabolism. Not surprisingly, the ketogenic diet induces a downregulation of the gene signature for cholesterol synthesis, as was also noted for the low protein diet.

Beside these similarities, I noticed some fundamental differences between the low protein and ketogenic diets. The ketogenic diet has a distinctive induction of the gene signature for the TFEB targets, a master regulator of lysosomal and autophagy related genes ^14^. The lysosomal pathway could be induced to mediate the endocytosis and degradation of the increased dietary fat. It could also be linked to the amino acid stress response, to mediate the degradation of proteins, but this is not observed in the context of a low protein diet (Fig. 2A). The ketogenic diet also induces a gene signature of adipogenesis (Fig. 1B), probably reflecting a lipid storage response.

The induction of the serine synthesis genes in the liver is corroborated by data from a second study (GSE115342 ^15^, Fig. 2C). The latter study also reported the gene expression profiles for the cerebral cortex, where two out of three samples exhibit an induction of the serine synthesis gene signature (Fig. 2D).

As a last case study, I consider a report of liver gene expression profiles of mice subject to a high fructose diet, high fat diet, dietary restricted and essential fatty acids restricted. These diets do not induce the serine synthesis gene signature across all samples (Fig. 2E). This evidence demonstrates that the ability of the low protein and ketogenic diets to induce the serine synthesis genes is specific to these diets and not a general response to dietary interventions.

### Pharmacological interventions, liver

Dietary restrictions while effective they may be hard to sustain. As an alternative we could think of pharmacological interventions. To identify compounds that induce the serine synthesis from 3-phosphoglycerate I took advantage of a toxicology study reporting the gene expression profiles of rats exposed to different compounds, including several drugs currently in use to treat a variety of illnesses ^16^. I have previously shown that this resource can be used to identify compounds that mimic calorie restriction ^8^. Here I will use the same approach to identify compounds that induces serine synthesis. The methodology is rather straightforward. First identify compounds that induce the serine synthesis gene signature in the 3 samples tested with a predefined dose and duration of the treatment. Then I extract those compounds that are positive for multiple doses/treatment-durations, using a hypergeometric test to quantify statistical significance.

The compounds at the top of the hit list target the oestrogen hormone pathway (Fig. 3A-E, Supplementary Table 2). That includes the non-steroidal oestrogen diethylstilbestrol, the oestrogen analogue ethynilstradiol, the aromatase inhibitor anastrazole and the oestrogen receptor modulator tamoxifen. In many instances the induction of the serine synthesis gene signature is matched by an induction of the ATF4 targets gene signature, suggesting that these compounds act through the amino acid stress response pathway. Whether that is the case remains to be validated.

**Figure 3.**
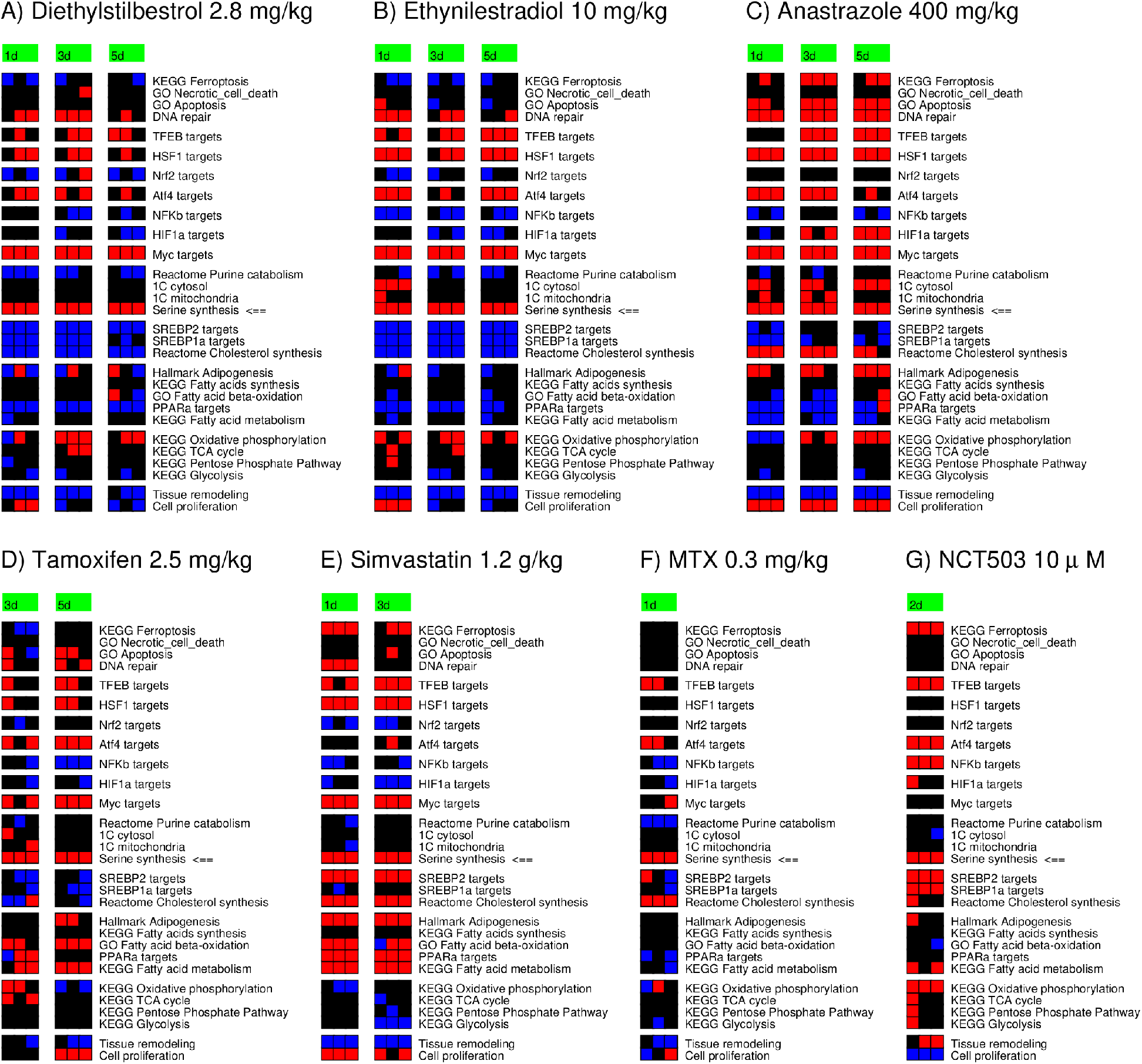
Pharmacological interventions inducing the expression of serine synthesis genes. A-F) Liver gene signatures in rats exposed to treatment with the indicated compounds. G) Gene signatures in a neuroblastoma cell line treated with the PHGDH inhibitor NCT503. Columns represent samples clustered by the treatment duration (in days) and rows represent gene signatures. <⩵ Points to the serine synthesis gene signature. Red indicates a significant gene signature induction relative to the average in controls, blue a significant repression and black no significant change.

Surprisingly Simvastatin, which targets the cholesterol synthesis pathway, also induces the serine synthesis gene signature (Fig. 3E). In this case there is no significant induction of the ATF4 targets gene signature, suggesting a different mechanism of action. It is worth noticing that the treatment with statin also induces the cholesterol synthesis gene signature (Fig 2E). I went back to the dataset an pulled down other statins tested, including atorvastatin, fluvastatin and lovastatin. These statins induce the serine synthesis gene signature as well, albeit not under all conditions tested (Supplementary Fig. 1). synthesis genes as a compensatory mechanism.

### Pharmacological interventions, hepatocytes

It is worth asking whether the same hits could have been identified using a cell culture system. The toxicology study in rats reported the gene expression of hepatocytes in culture treated with a similar collection of compounds (GSE57805 ^16^). Although I found no significant hit when considering the hepatocytes gene expression profiles, there are several compounds with a border line significance (~0.07, Supplementary Table 2). That list contains 27 compounds, including the oestrogen analogue ethinylestradiol and some statins, but missing diethylstilbestrol, anastrozole and tamoxifen. These mismatches highlight the requirement to conduct the investigation in whole organisms.

### Pharmacological interventions, other tissues

The toxicology study in rats reported the gene expression profiles of heart, kidney and thigh muscle of mice treated with different selections of compounds ^16^. Among the compounds tested, I did not find any compound reaching statistical significance for the induction of serine synthesis genes. However, it should be noted that the compounds inducing the serine synthesis genes in the liver were not tested in the heart and thigh muscle datasets. At this point I can only conclude that they do not induce the serine synthesis genes in the kidney.

## Discussion

I have identified pharmacological interventions increasing the liver expression of genes in the pathway of serine synthesis from the glycolytic intermediate 3-phosphoglycerate. Most convincing is the action of compounds modulating the oestrogen hormone pathway and statins, as demonstrated by the induction of serine synthesis genes for several compounds on those classes. With less confidence due to the lack of sufficient testing, I identified the antifolate methotrexate and a serine synthesis inhibitor as additional pharmacological interventions with the potential to induce the serine synthesis genes in the liver. In all cases the induction of serine synthesis genes could be a feedback mechanism to increase the availability of serine, glycine or formate.

I have presented evidence for the pharmacological induction of serine synthesis genes in the liver. The compounds testing positive for induction of serine synthesis genes in the liver (diethylstilbestrol, ethynylstradiol, anastrazole, tamoxifen, simvastatin) were not included in the heart and thigh tissues gene expression datasets. What happens in other tissues requires further investigation.

I have uncovered that a low protein or a ketogenic diet can induce the expression of serine synthesis genes in the liver relative to a control diet. The low protein diet recapitulates the induction of the amino acid stress response observed in cell culture systems. The ketogenic diet case is a reminder that glucose are not empty calories. The induction of serine synthesis genes in the liver by a ketogenic diet is probably another feedback mechanism. The reduced availability of glucose and as a consequence the glycolytic intermediate 3-phosphoglycerate, the substrate for serine synthesis. The system would them compensate by inducing the serine synthesis genes.

It does not scape to my attention that the ketogenic diet was first introduced for the management of epilepsy ^19^. D-serine and glycine, two downstream products of serine metabolism (Fig. 1) are co-agonists of NMDA receptor. D-serine and glycine potentiate the activity of anticonvulsant drugs in animal models ^21,22^. Pharmacological inhibition of PHGDH, the first enzyme in serine synthesis, reduces the production of D-serine and glycine in cultures of astrocytes and brain hippocampus slices and inhibits the NMDA receptor response ^20^. Here I have shown that a ketogenic diet induces the expression of serine synthesis genes in the liver and the cerebral cortex of mice. In the brain the proteinogenic amino acid L-serine is converted to D-serine and broken down to glycine and formate as well ^6^. These observations point to the induction of serine synthesis genes as the mechanism of action of a ketogenic diet in the management of epilepsy.

Beyond epilepsy, dietary and pharmacological induction of serine synthesis from 3-phosphoglycerate should be investigated for the management of other neurological disorders of serine deficiency ^23^. The same applies to macular telangiectasia type 2 and peripheral neuropathies where serine deficiency has been linked to significant changes in the phospholipids profile ^2^.

Finally, I would like to bring to your attention that serine synthesis from the glycolytic intermediate 3-phosphoglycerate is effectively a sink for glycolysis and gluconeogenesis (Fig. 1). The induction of serine synthesis could be an effective strategy to lower glucose levels for the management of hyperglycaemia in diabetes and obese patients.

## Acknowledgements

This work was supported by Cancer Research UK C596/A21140.

## Author contributions

AV conceived the work and wrote the manuscript.

## Data availability

All the gene expression data analysed in this work is available from Gene Expression Omnibus.

---

**Supplementary Figure 1.**
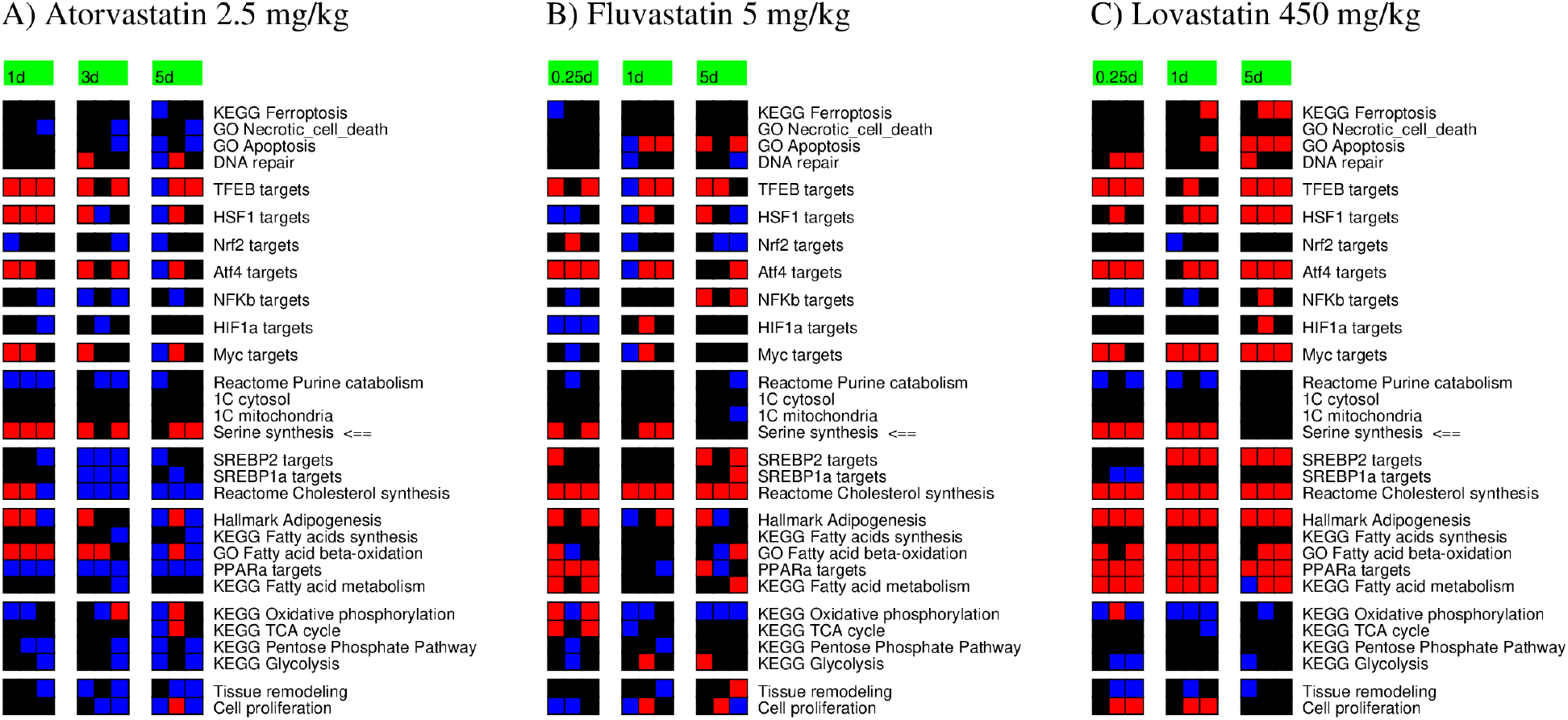
Liver gene signatures following treatment with statins. Columns represent samples clustered by the treatment duration (in days) and rows represent gene signatures. <⩵ Points to the serine synthesis gene signature. Red indicates a significant gene signature induction relative to the average in controls, blue a significant repression and black no significant change.

